# Discovery of human ACE2 variants with altered recognition by the SARS-CoV-2 spike protein

**DOI:** 10.1101/2020.09.17.301861

**Authors:** Pete Heinzelman, Philip A. Romero

## Abstract

Understanding how human ACE2 genetic variants differ in their recognition by SARS-CoV-2 can have a major impact in leveraging ACE2 as an axis for treating and preventing COVID-19. In this work, we experimentally interrogate thousands of ACE2 mutants to identify over one hundred human single-nucleotide variants (SNVs) that are likely to have altered recognition by the virus, and make the complementary discovery that ACE2 residues distant from the spike interface can have a strong influence upon the ACE2-spike interaction. These findings illuminate new links between ACE2 sequence and spike recognition, and will find wide-ranging utility in SARS-CoV-2 fundamental research, epidemiological analyses, and clinical trial design.

## Main

The highly contagious and pathogenic SARS-CoV-2 coronavirus has rapidly spread worldwide, leading to a global public health emergency. The virus recognizes and infects human cells by binding to the angiotensin-converting enzyme 2 (ACE2) protein, which is expressed on the surface of epithelial cells in human lungs. The high-affinity interaction between the virus and ACE2 is a major determinant of SARS-CoV-2’s high infectivity [1,2]. Characterizing how this interaction varies across the human population is of great importance for biomedical research, clinical trial design, and retrospective analysis of epidemiological data to facilitate development of vaccines, therapeutics, and treatment regimens that mitigate or prevent the acute and prolonged effects of COVID-19.

In this work, we study how missense single-nucleotide variants (SNVs) of the ACE2 protein influence recognition by the SARS-CoV-2 spike protein. In particular, we perform deep mutational scanning (DMS) to experimentally map how over 3,500 amino acid substitutions in ACE2’s extracellular peptidase domain impact binding to spike. In addition to capturing known residues that are key to spike binding and revealing dozens of new ACE2 sites that influence the ACE2-spike interaction, our DMS data identified over 100 human SNVs that are likely to exhibit altered spike binding. These new insights will have an important impact toward understanding and combating SARS-CoV-2 infection.

We performed a deep mutational scan on ACE2’s extracellular peptidase domain (codons 18-615) using a combination of random mutagenesis, yeast surface display (YSD)-based screening, and next-generation sequencing. We used error-prone PCR to randomly mutagenize the ACE2 gene, and cloned the resultant library into a yeast surface display (YSD) vector [3] that anchors the ACE-2 C-terminus to the yeast cell wall (Supplementary Figure 1). ACE2-displaying yeast were incubated with a fluorescently-labeled SARS-CoV-2 spike receptor binding domain (RBD) and fluorescence activated cell sorting (FACS) was used to enrich RBD-binding ACE2 variants (Supplementary Figures 2 & 3). We then performed Illumina sequencing of both the initial ACE2 library and enriched ACE2 variants, and analyzed the resulting data to assess how individual amino acid substitutions affect spike protein binding.

The data produces a SARS-CoV-2 spike-binding map for 3571 amino acid substitutions across 597 positions in ACE2’s peptidase domain (Fig. 1ab). 68% of these substitutions decrease spike binding, 4% increase spike binding, and 28% have no statistically significant effect. Importantly, this set of observed amino acid substitutions contains 165 of the 196 (84%) annotated ACE2 missense SNVs found in the human population [4].

**Figure 1:**
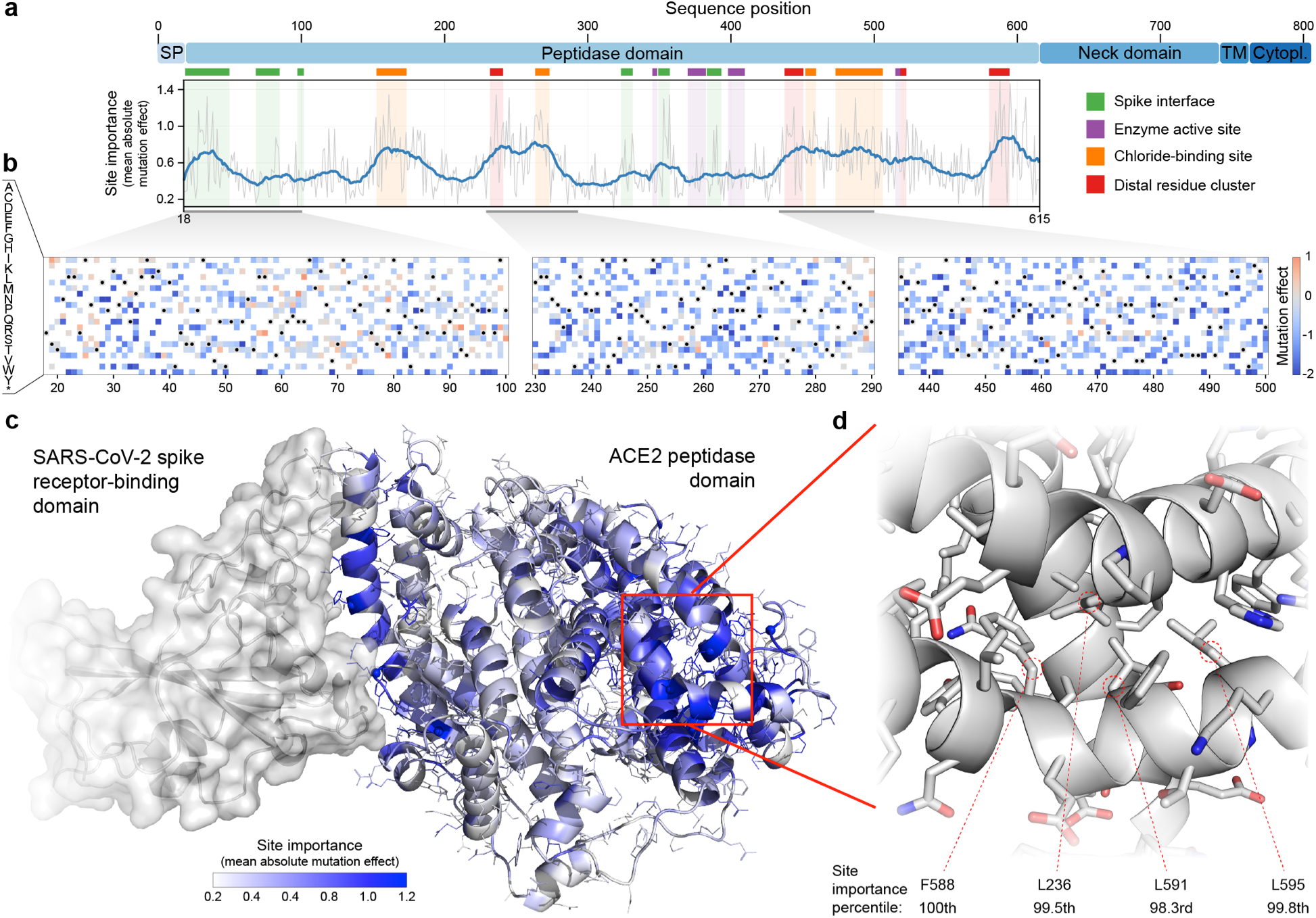
Large-scale mutagenesis of ACE2’s peptidase domain. (a) Analysis of how amino acid substitutions across positions 18-615 affect binding of the SARS-CoV-2 spike protein. The plot quantifies the importance of each site by taking the mean of the absolute value of all mutation effects observed at that site. The grey line represents the mean absolute value of the mutation effect and the blue line shows the moving average to highlight general regions of ACE2 that are important for binding. Key structural landmarks are highlighted with shaded regions along the length of the sequence. (b) Mutation effect heat maps for three different regions of ACE2. Red denotes mutations that increase ACE2 spike binding; blue denotes reduced binding. Overall, we observe the effects of 3571 amino acid substitutions across 597 positions in ACE2’s peptidase domain. (c) The mean absolute mutation effect mapped onto the three-dimensional ACE2 structure (PDB ID: 6LZG). Residues near the spike interface are important for binding, in addition to many sites located on the distal lobe of the protein domain. (d) The most important region of ACE2 structure is composed of a tightly packed cluster of residues located over 30 Å from the spike interface.

Our DMS data provides a comprehensive view of how each ACE2 site influences spike binding (Fig 1). Previous work has focused on the interface between the RBD and ACE2’s N-terminal alpha helices and a short beta turn present in the middle of the ACE2 sequence [5]. Known key ACE2 residues at this binding interface include S19, Q24, D30, H34, D38, Y41, Q42, Y83, and K353. We find that most of these residues are crucial for spike binding in our DMS experiment, ranking above the 90^th^ percentile for site importance as quantified by the mean absolute mutation effect (Fig 1). This agreement with known structural interactions validates the quality of our DMS data.

Our DMS data also revealed previously undescribed relationships between ACE2 and spike binding. Residues within the ACE2 peptidase active site have little influence on spike protein binding, which is expected since ACE2’s natural peptidase function is distinct from viral recognition. Surprisingly, we found residues near ACE2’s chloride-binding site, which is important in regulating ACE2 peptidase activity [6], play an important role in spike binding. This site is over 40 Å from the spike interface and thus mutations contained therein may influence spike binding by changing ACE2’s conformation. Our analysis also identified a tightly packed cluster of residues that strongly influence spike binding (Fig 1d). Residues L236, F588, L591, and L595 are ranked above the 98^th^ percentile for site importance despite being over 30 Å from the spike interface. As with chloride binding site mutations, substitutions at this distal residue cluster may impact spike binding by altering ACE2’s conformation.

Our DMS data allows assignment of putative spike-binding annotations to 165 missense SNVs (Fig 2a). 68% of these missense SNVs decrease spike binding, 27% bind similar to wild-type ACE2, and 5% possess increased binding. A vast majority of the SNVs that alter spike binding are distant from the spike interface that has been the focus of prior ACE2 mutagenesis studies [1,2,7].

**Figure 2:**
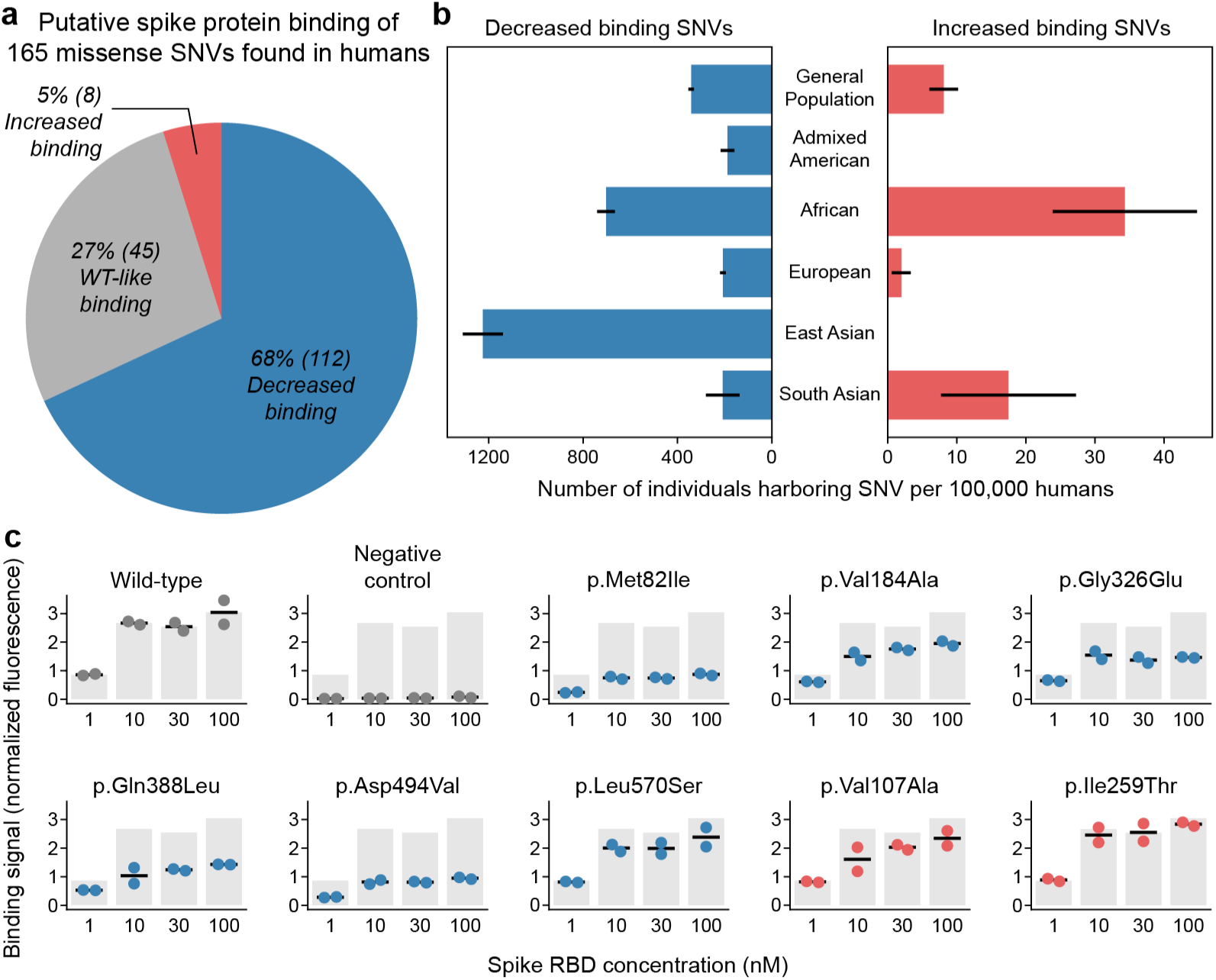
Functional annotation of human missense SNVs. (a) Our large-scale mutagenesis data encompasses 165 missense SNVs found in the human population. We assign putative spike protein binding annotations to these variants. Roughly two-thirds display decreased spike binding, one-third display binding similar to wild-type ACE2, and a small fraction display increased binding. (b) We estimate 320-365 individuals per 100,000 humans may harbor SNVs that decrease spike binding, while 4-12 individuals may have SNVs that increase binding. Specific human subpopulations possess higher and lower frequencies of SNVs that alter spike binding. Error bars represent one standard deviation and were generated by applying bootstrap statistical analysis to the SNV allele count data. (c) Data for YSD-based binding characterization of eight missense SNVs. The binding signal is defined as mean fluorescence for yeast-displayed ACE2 spike binding divided by mean fluorescence corresponding to ACE2 display level on the yeast surface (Supplementary Figures 4 & 5). This normalization is applied to prevent potential differences in ACE2 display from biasing assessment of spike binding. SNVs with negative mutation effect values appear as blue dots, while SNVs with positive mutation effects are shown in red. Each dot represents an experimental replicate at varying spike protein concentrations. The grey bars show wild-type ACE2’s binding signal to serve as a reference.

Based on each variant’s allele frequency in the general population, we estimate that 320-365 (95% confidence interval) individuals per 100,000 humans possess an SNV that may decrease spike protein binding, while an estimated 4-12 (95% confidence interval) of every 100,000 persons possess SNVs that may increase spike binding (Fig 2b). The most highly observed SNVs appear at frequencies approaching 1-in-1000 humans (Supplementary Table 1), indicating that millions of individuals may possess these SNVs, and speaks to the feasibility of epidemiological studies and clinical investigations focused on human genetic variants that influence the ACE2-spike interaction (Supplementary Table 1). In addition, certain SNVs are more prevalent among individuals of different ancestries, further supporting the feasibility of assembling large groups of ACE2 isogenic human subjects. The frequency of individuals harboring an SNV that alters spike binding may vary by over six-fold between different ancestries (Fig 2b), and frequencies of specific SNVs, such as p.Met82Ile in the African population, are increased by more than a factor of five relative to the general human population (Supplementary Table 1).

We chose ten missense SNVs for further characterization based on their high representation in the human population and the magnitudes of their mutation effects as calculated from the DMS data (Supplementary Table 2). We characterized these ten variants using our YSD flow cytometry binding assay at varying concentrations of spike protein. Six of eight putative decreased binding SNVs possessed markedly reduced spike binding relative to wild type ACE2, while the two putative increased binding SNVs returned spike binding signal values similar to or less than wild type (Fig 2c). Notably, p.Asp494Val substantially reduced spike binding, but has not been examined in computational work focused on how SNVs impact the ACE2-spike interaction [8,9], and was excluded from site-directed ACE2 mutagenesis studies [2,7]. This result highlights the ability of DMS to discover new SNVs that are as important as interface residues for binding to the spike protein.

The spike titration data gives insight into the mechanism of each SNV’s altered spike binding. Wild-type ACE2 exhibits a step-like response between 1 and 10 nM; such behavior is expected for standard sigmoidal binding curves as the antigen concentration crosses the *K*_*D*_ value. The binding curves for many of the variants with decreased binding also exhibit a step-like response similar to that for wild type ACE2, but possess maximum binding signals that are reduced by up to three-fold. This reduction in maximum binding signal may be the result of the proteins existing in a mixture of conformational states, all or some of which feature higher *K*_*D*_ values than wild

type ACE2 on the yeast surface. Previous yeast display studies of human brain-derived neurotrophic factor have observed single protein variants that exist in multiple conformational states, each of which possess their own unique *K*_*D*_ values [10].

Two of the tested SNVs with large negative mutation effect values, p.Ala242Val and p.Tyr252Cys, exhibited reduced display on the yeast surface and inability to bind spike protein (Supplementary Figure 6). This may be due to destabilization of the protein structure on the yeast surface or aberrant processing in the yeast protein secretory pathway [11,12].

The ability of SARS-CoV-2 virions to enter and infect human cells is dependent upon a number of ACE2 properties: the binding affinity for the SARS-CoV-2 spike, the amount of ACE2 that migrates through the endoplasmic reticulum to reach the cell surface, and the turnover rate of ACE2 at the cell membrane [13]. Our work has identified ACE2 SNVs that are markedly distinct from the wild-type protein with respect to the above properties, and thus present new opportunities to interleave ACE2 biochemical studies, ACE2 variant animal experiments, and observations of the human population. This work has also revealed that mutations in ACE2 residues distal to the SARS-CoV-2 spike-binding interface may alter ACE2 properties relevant to SARS-CoV-2 recognition; this finding will motivate future empirical and *in silico* research efforts to expand their scope beyond the spike binding interface. Collectively our results provide key insights that can be readily actioned toward fundamental research, clinical studies, and epidemiological analyses for the treatment and prevention of SARS-CoV-2 infection.

## Methods

### ACE2 library generation, screening, and sequencing

Residues 18-165 of the human ACE2 gene were synthesized as a yeast codon-optimized gBlock (Integrated DNA Technologies, Coralville, IA) and the gBlock was ligated into the unique NheI and MluI sites of the yeast surface display vector VLRB.2D-aga2 (provided by Dane Wittrup, MIT); this yeast display vector fuses the aga2 protein to the C-terminus of ACE2 (Supplementary Figure 1). The ACE2 gene contained His to Asn mutations at positions 376 and 380 to abolish zinc binding and ACE2 proteolytic activity. The GeneMorph II Kit (Agilent Technologies, Santa Clara, CA) was employed to generate a human ACE2 random mutant library using the wild type ACE2 display plasmid as template and respective forward and reverse primers CDspLt (5’-GTCTTGTTGGCTATCTTCGCTG-3’) and CDspRt (5’-GTCGTTGACAAAGAGTACG-3’). Error prone PCR products from the GeneMorph random mutagenesis reactions were digested NheI to MluI and ligated into the VLRB.2D-aga2 vector digested with these same enzymes. Ligation products were concentrated and desalted using the Zymoclean Clean & Concentrator 5 kit (Zymo Research, Orange, CA) and electroporated into 10G Supreme *E. coli* (Lucigen, Middleton, WI). Transformants were pooled and cultured in LB media containing 100ug/mL carbenecillin overnight at 30°C and plasmids subsequently harvested using the Qiagen (Valencia, CA) Spin Miniprep kit.

Yeast display *Saccharomyces cerevisiae* strain EBY100 was made competent using the Sigma-Aldrich yeast transformation kit. Transformants were pooled and cultured in low-pH Sabouraud Dextrose Casamino Acid media (SDCAA, per liter - 20 g dextrose, 6.7 grams yeast nitrogen base (VWR Scientific, Radnor, PA), 5 g Casamino Acids (VWR), Citrate buffer (pH 4.5) - 10.4 g sodium citrate / 7.4 g citric acid monohydrate) at 30°C and 250 rpm for two days. For induction of ACE2 mutant library display a 5 mL Sabouraud Galactose Casamino Acid (SGCAA, Per liter - Phosphate buffer (pH 7.4) - 8.6 g NaH_2_PO*H_2_O / 5.4 g Na_2_HPO_4_, 20 g galactose, 6.7 g yeast nitrogen base, 5 g Casamino Acids) culture was started at an optical density, as measured at 600 nm, of 0.5 and shaken overnight at 250 rpm and 20°C.

After overnight induction approximately 3*10^6^ yeast cells were harvested by centrifugation, washed once in pH 7.4 Phosphate Buffered Saline (PBS) containing 0.2% (w/v) bovine serum albumin (BSA) and incubated overnight in 500 µL of PBS/0.2% BSA containing 150 nM His_6_-tagged SARS-CoV-2 spike RBD (Sino Biological, Chesterbrook, PA) and 5 µg/mL anti-*myc* IgY (Aves Labs, Tigard, OR) at 4°C on a tube rotator at 18 rpm. Incubation with a concentration of spike, i.e., 150 nM, well above the ACE2-spike RBD equilibrium binding dissociation constant (*K*_*D*_) value was employed to maximize depletion of ACE2 mutants with very low or no binding to the spike RBD during FACS. Following overnight incubation yeast were washed once in PBS/0.2% BSA and rotated at 18 rpm in same buffer containing 5 µg/mL mouse anti-His_6_ IgG (BioLegend, San Diego, CA) that had been conjugated with Alexa647 using the N-hydroxysuccinimidyl ester (Molecular Probes, Eugene, OR) of this fluorophore and 2 µg/mL Alexa488-conjugated goat anti-chicken IgG (Jackson ImmunoResearch, West Grove, PA) for one hour at 4°C. Yeast cells were subsequently washed and resuspended in ice cold PBS for sorting on a FACS Aria III (Becton Dickinson, Franklin Lakes, NJ) located in the University of Wisconsin-Madison Carbone Cancer Center Flow Cytometry Laboratory. Sorting gates were set such that ACE2 mutant library members at or above approximately the 80^th^ percentile with respect to binding to the CoV-2 spike RBD were isolated (Supplementary Figures 2 & 3).

Yeast cells isolated during sorting were cultured overnight in low pH SDCAA media at 30°C with shaking at 250 rpm and the following morning 1/10^th^ of the culture volume was expanded in low pH SDCAA to an OD of 0.1, shaken at 30°C and 250 rpm, and harvested by centrifugation after the OD had reached 0.4. ACE2 mutant library yeast display plasmids were rescued using the ZymoPrep Yeast Plasmid Miniprep II kit (Zymo Research). Rescued plasmids were amplified via electroporation into 10G Supreme *E. coli* with subsequent culturing and harvesting as described above. For NGS analysis, amplified plasmid DNA from the post-FACS yeast population and unsorted ACE2 random mutant library yeast display plasmid DNA were digested in separate reactions using PstI and XhoI restriction enzymes. Digested plasmids were run in a 1.2% agarose gel and the ACE2 band was excised and purified using the Zymo Gel Extraction kit (Zymo Research). Purified DNA was submitted to the University of Wisconsin-Madison Biotechnology Center DNA Sequencing Facility for analysis.

To assess the effects of the respective ACE2 missense mutations noted in Figure 2 on CoV-2 spike binding affinity, these mutations were introduced into the wild type ACE2 gene via overlap extension PCR and resultant mutant genes were ligated into the VLRB.2D-aga2 yeast display vector using NheI and MluI sites as described above. Ligations were subsequently transformed into NEB-5α chemically competent *E. coli* (New England Biolabs, Beverly, MA) and single colonies picked for culturing, plasmid harvest, and sequencing to verify correct introduction of SNV mutations. Plasmid DNA for each respective member of the collection of SNV ACE2 genes was transformed into EBY100 yeast made competent using the Zymo Research Frozen EZ Yeast Transformation II kit with transformants grown on synthetic dropout (SD) -Trp (MP Biomedicals, Irvine, CA) agar plates for two days at 30°C.

After two days of growth on SD -Trp agar plates respective yeast colonies for SNV and wild type ACE2 yeast were picked into 4 mL of low pH SDCAA and grown overnight at 30°C with shaking at 250 rpm. These cultures were subsequently induced in 5 mL of SGCAA overnight at 20°C with shaking at 250 rpm; induction culture starting OD was 0.5. After overnight induction of ACE2 surface display yeast cells for each ACE2 variant to be analyzed were harvested by centrifugation and washed once as described above. 2*10^5^ yeast cells were tumbled overnight at 4°C in 300 µL of PBS/0.2% BSA containing 5 µg/mL anti-*myc* IgY and various concentrations of His_6_-tagged CoV-2 spike RBD as denoted in Figure 2. Following secondary labeling with fluorescent anti-His_6_ and anti-IgY antibodies yeast were washed once with PBS/0.2% BSA and resuspended in ice cold PBS for flow cytometric analysis. Analyses were performed using a Fortessa analyzer (Becton Dickinson) in the University of Wisconsin-Madison Biochemical Sciences Building.

### Analysis of deep mutational scanning data

The deep mutational scan generated Illumina sequencing data for the unsorted and sorted ACE2 libraries. Paired-end reads were mapped to the ACE2 gene using Bowtie2’s very-sensitive-local alignment setting [14]. The resulting SAM files were filtered to remove sites with a Phred quality score (Q score) of less than 30 and translated to amino acid sequences. The entire data set was filtered to remove amino acid substitutions with less than 10 observations.

The unsorted and sorted ACE2 amino acid sequences were then analyzed using a positive-unlabeled (PU) learning method that estimates how amino acid substitutions affect a protein’s function [15]. The PU learning method returns an estimated coefficient, also referred to as the mutation effect, and a p-value corresponding to each amino acid substitution. Coefficients with a p-value less than 0.05 were considered to have a significant effect on spike protein binding. The importance of each site in ACE2 was evaluated by aggregating all mutation effects at a given position. We calculated the mean absolute value of all coefficients at each site to generate a “site importance” profile across the extracellular domain of ACE2. A site with a large mean absolute mutation effect is likely to be important for spike protein binding.

### Analysis of human genetic data

Human genetic data was retrieved from the Genome Aggregation Database (gnomAD) [4]. Exome data from gnomAD v2.1.1 and genome data from gnomAD v3 were combined to generate a data set representing over 250,000 individuals. Allele frequencies for each missense single-nucleotide variant (SNV) were calculated for the general population (all samples), and Admixed American (Latino), African, European, East Asian, and South Asian subpopulations. The European subpopulation combined Ashkenazi Jewish, Finnish, and non-Finnish European samples because none of these sub-subpopulations had a high frequency of SNVs that alter spike binding.

Confidence intervals for the fraction of individuals harboring an SNV that alters spike binding were calculated by bootstrapping the allele count data. For each SNV in each subpopulation, a binomial random variable was generated using a proportion equal to the observed allele frequency and a number of trials equal to the observed allele number. This binomial random variable was used to calculate a resampled allele frequency, and the frequency of SNVs at each site was combined to estimate the fraction of individuals harboring any SNV that alters spike binding. This resampling procedure was repeated 1,000 times to obtain a distribution over these estimates.

## Acknowledgements

This work was supported by the US National Institutes of Health (5R35GM119854).

## Author contributions

P.H. and P.A.R conceived the project. P.H. performed the experiments. P.H. and P.A.R analyzed the data. P.H. and P.A.R. wrote the manuscript.

