## Supplementary Figures for "Discovery of human ACE2 variants with altered recognition by the SARS-CoV-2 spike protein"

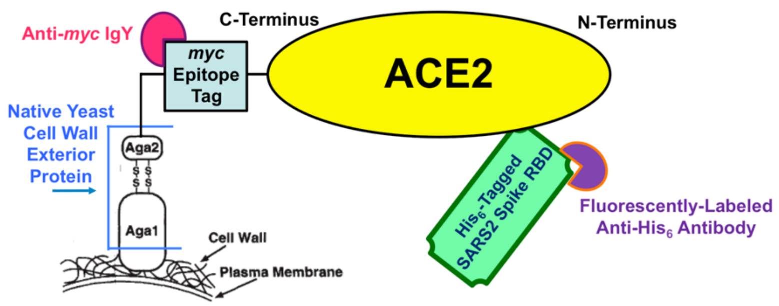


**Supplementary Figure 1.** Yeast display schematic. ACE2's C-terminus was fused to a Myc epitope tag and the Aga2 native yeast surface protein on yeast cell wall. ACE2 display was quantified using an anti-*myc* chicken IgY that is detected using an Alexa488-conjugated anti-chicken goat IgG as secondary label (not depicted). Binding of yeast-displayed ACE2 to $His_6$-tagged SARS-CoV-2 spike RBD was detected via incubation with Alexa647-conjugated anti-$His_6$ mouse IgG. Both ACE2 display and ACE2 binding to spike RBD were measured by flow cytometry. Each yeast cell displays up to $10^4$ copies of a single ACE2 variant on its surface.

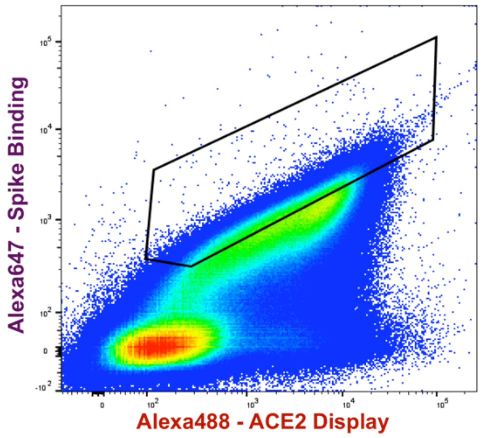


**Supplementary Figure 2.** Dot plot for FACS of random mutant ACE2 library. Sort gate (black polygon) was defined to enrich the top twenty percent of *myc*-positive (ACE2 displaying) yeast. The X-axis denotes Alexa488 fluorescence (ACE2 display) and the Y-axis denotes Alexa647 fluorescence (ACE2 binding to spike protein). The plot depicts dots for approximately $1.5*10^6$ yeast cells. The library was incubated with 150 nM spike RBD prior to sorting. The high density (red) oval at the lower left primarily contains yeast that have not been induced. For biological reasons that are poorly understood, even homogeneous populations of yeast carrying identical display plasmids, i.e., wild type ACE2, feature 25% or greater cells that do not display any protein.

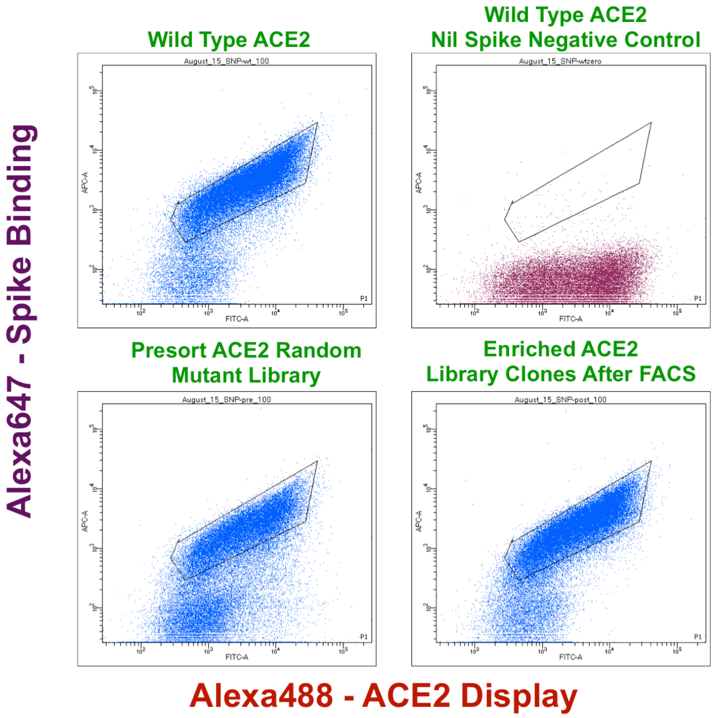


**Supplementary Figure 3.** Flow cytometry dot plots for ACE2 random mutant library before and after FACS enrichment. The X-axes denote Alexa488 fluorescence (ACE2 display) and the Y-axes denote Alexa647 fluorescence (ACE2 binding to spike protein). The plots depict dots for approximately $5*10^4$ yeast cells. The yeast were incubated with 100 nM spike RBD, or no spike for the wild-type ACE2 nil spike negative control sample, prior to flow cytometric analysis. For biological reasons that are poorly understood, even homogeneous populations of yeast carrying identical display plasmids, i.e., wild-type ACE2, feature 25% or greater cells (lower left region of plots) that do not display any protein.

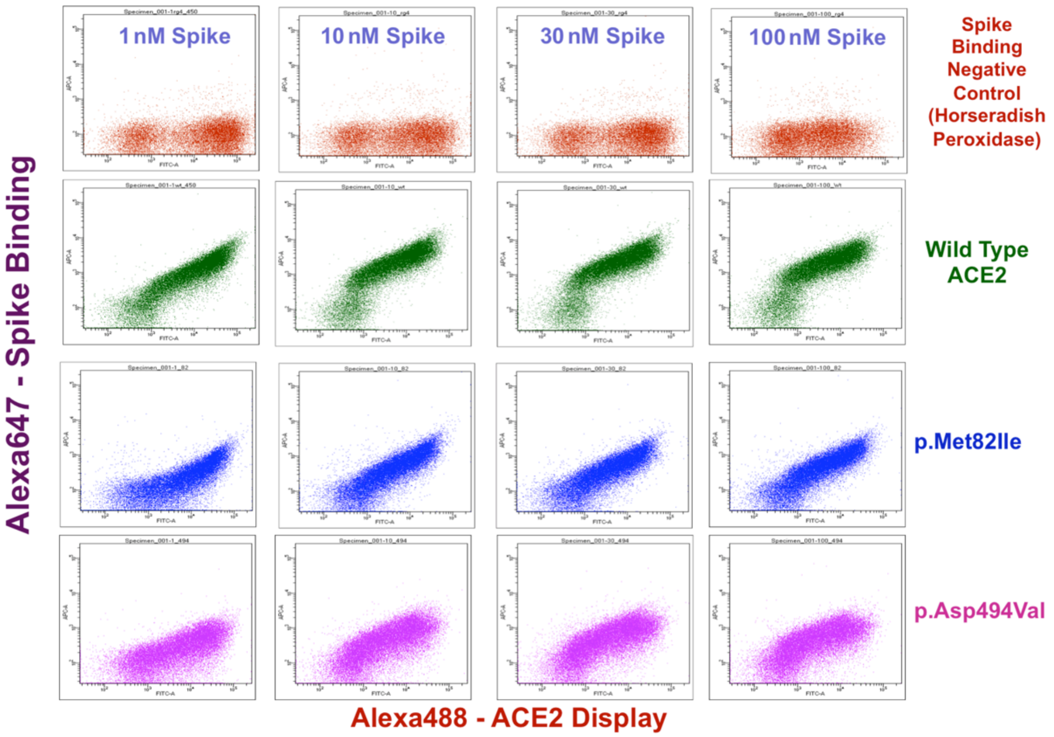


**Supplementary Figure 4.** Representative flow cytometry dot plots corresponding to binding signal results presented in Figure 2. The X-axes denote Alexa488 fluorescence (ACE2 display) and the Y-axes denote Alexa647 fluorescence (ACE2 binding to spike protein). The plots depict dots for approximately $3*10^4$ yeast cells. The yeast were incubated with spike RBD at concentrations noted in figure prior to analysis. For biological reasons that are poorly understood, even homogeneous populations of yeast carrying identical display plasmids, i.e., wild-type ACE2, feature 25% or greater cells (lower left region of plots) that do not display any protein.

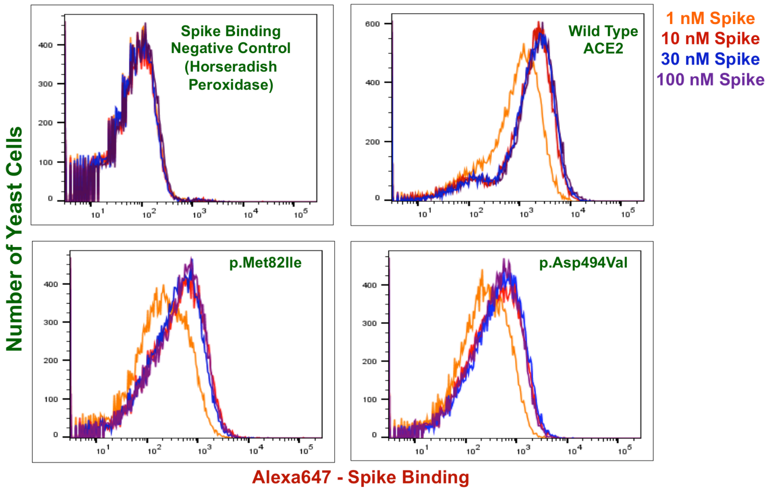


**Supplementary Figure 5.** Representative flow cytometry histogram overlays corresponding to results presented in Figure 2. The X-axes denote Alexa647 fluorescence (ACE2 binding to spike protein), and the Y-axes denote the number of yeast cells. Each histogram represents approximately $3*10^4$ yeast cells. The yeast were incubated with spike RBD at concentrations noted in the figure prior to analysis. For biological reasons that are poorly understood, even homogeneous populations of yeast carrying identical display plasmids, i.e., wild-type ACE2, feature 25% or greater cells (leftmost yeast in ACE2 histograms) that do not display any protein.

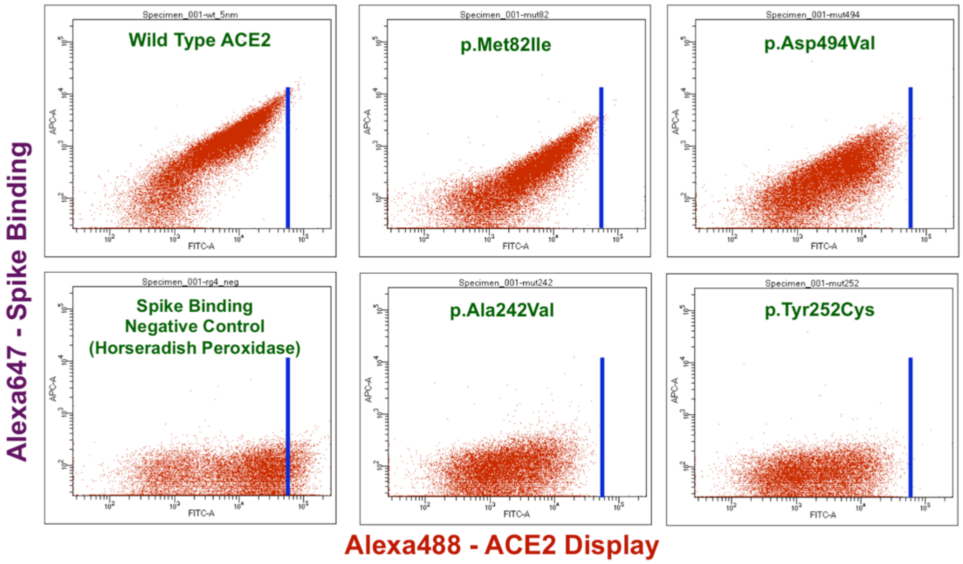


**Supplementary Figure 6.** Flow cytometry dot plots for wild type and selected SNV ACE2s. The X-axes denote Alexa488 fluorescence (ACE2 display) and Y-axes denote Alexa647 fluorescence (ACE2 binding to spike protein). The plots depict dots for approximately $3*10^4$ yeast cells. The dark blue line denotes upper limit of Alexa488 signal (ACE2 display) for wild-type ACE2. The yeast were incubated with 5 nM spike RBD prior to flow cytometric analysis. For biological reasons that are poorly understood, even homogeneous populations of yeast carrying identical display plasmids, i.e., wild-type ACE2, feature 25% or greater cells (lower left region of plots) that do not display any protein.
